# Insular cortex encodes task alignment

**DOI:** 10.1101/2024.12.13.628250

**Authors:** Johannes Niediek, Maciej M. Jankowski, Ana Polterovich, Alexander Kazakov, Israel Nelken

## Abstract

Animals can learn complex behaviors. Animal behavior in the lab has traditionally been studied via summary statistics such as trial-based success rates. However, animal behavior is much more fine-grained: a trial in an experiment often consists of multiple actions, and more than one strategy can lead to a successful completion of a trial. To understand how the brain controls behavior, a fine-grained yet compact description of behavior is necessary.

We describe here an approach for estimating the strategies animals use for a large family of tasks from first principles. Using reinforcement learning with informational constraints on policies, we compute a rich set of candidate policies with a small number of meaningful parameters, and match observed behavior to these policies. In a sample rat task, our approach revealed ongoing learning for more than 100 days after the saturation of success rates. Moreover, we showed that many neurons in the insular cortex of rats track the instantaneous task engagement of the rats with a resolution of a few minutes.

Due to its generic formulation in reinforcement learning terminology, our work is directly applicable to the majority of animal tasks in use today.

## Introduction

The behavior of animals unfolds in time, and success in any behavioral task requires multiple actions that may span many seconds. Fortunately, recent technology enables the tracking of animal behavior at unprecedented spatial and temporal resolutions (e.g., Ibáñez Alcalá et al. 2024; Nourizonoz et al. 2020; Ballesta et al. 2014). The availability of such high-resolution behavioral data calls for novel approaches to its analysis.

Common descriptions of behavior often summarize many seconds of behavior by a single number (e.g., trial-based success probability). However, trials usually consist of multiple actions, and in a complex task, multiple strategies can lead to the successful completion of a trial. Current approaches to modeling behavioral strategies in relation to a task include a mixture-of-agents approach (Miller, Botvinick, and Brody 2017), Hidden Markov Models (Ashwood et al. 2022), a combination of both (Venditto et al. 2024), or a reinforcement-learning based action-propensity model (Kastner et al. 2022). Other approaches use hidden states or contexts (e.g. Heald, Lengyel, and Wolpert 2021). However, these approaches aim at modeling animal behavior in a coarse-grained way, by focusing on a few decisions per trial, underplaying the richness of the observed behavior that is apparent when considering fine-grained actions at the sub-trial level.

Here, we propose a novel, theoretically grounded approach for the analysis of animal behavior in complex tasks that takes into account animal actions at a sub-trial, sub-second level. All states are fully observable, and the model can accommodate many actions in each state and action sequences of arbitrary length per trial. The task performed by the animal is modeled as a Markov Decision Process (MDP; Sutton and Barto 2018). The central notion for describing agent behavior in an MDP is that of a “policy” – a function that selects the actions to perform in each state of the process. Policies are notoriously hard to estimate from experimental data. The main contribution of this paper is a first-principles approach to compute a rich pool of policies of varying qualities, interpolating between task-agnostic policies and optimal policies. This pool of policies can be easily searched for a good match to the observed animal behavior, providing deep insights into the animal’s learning process of the task as well as into the relations with neural activity.

We evaluate our method on a complex task performed by rats in a novel, large behavioral arena (Jankowski, Polterovich, et al. 2023). The position of the best fitting policy in the range of policy qualities serves as a measure of task performance, which we call “task alignment”. We use task alignment to study rat performance in the task over time scales from minutes to many months. We show that task alignment continues to increase for many weeks past the saturation of the reward rate. At the same time, task alignment fluctuates at time scales of a few minutes during behavioral sessions. We argue that at this time scale, task alignment is really a measure of task engagement, and we show that firing rates of many neurons in the insular cortex strongly correlate with task alignment on a minute by minute basis.

## Results

### Rats quickly learned a complex task

To measure rat behavior at a high spatial and temporal resolution, we used the RIFF (Jankowski, Polterovich, et al. 2023), a large arena for jointly studying behavior and brain activity in freely moving rats (Fig. 1A). The RIFF is equipped with six interaction areas (IAs), each consisting of a food port and a water port together with nose-poke detectors and air-puff valves (Fig. 1A). Five female rats were trained in the RIFF. A schematic of the task is shown in Fig. 1B. In short, rats foraged in the arena, and had to correctly time their movements in order to receive reward (food pellets or water, at their preference). Three different strategies were available to the rats. Each strategy required the rat to move away from the last port of reward in different directions. Each training session in the RIFF lasted about 70 minutes, and sessions were performed daily on five consecutive days per week (also see Methods). Rats could freely choose their actions and strategies at any moment.

**Fig. 1.**
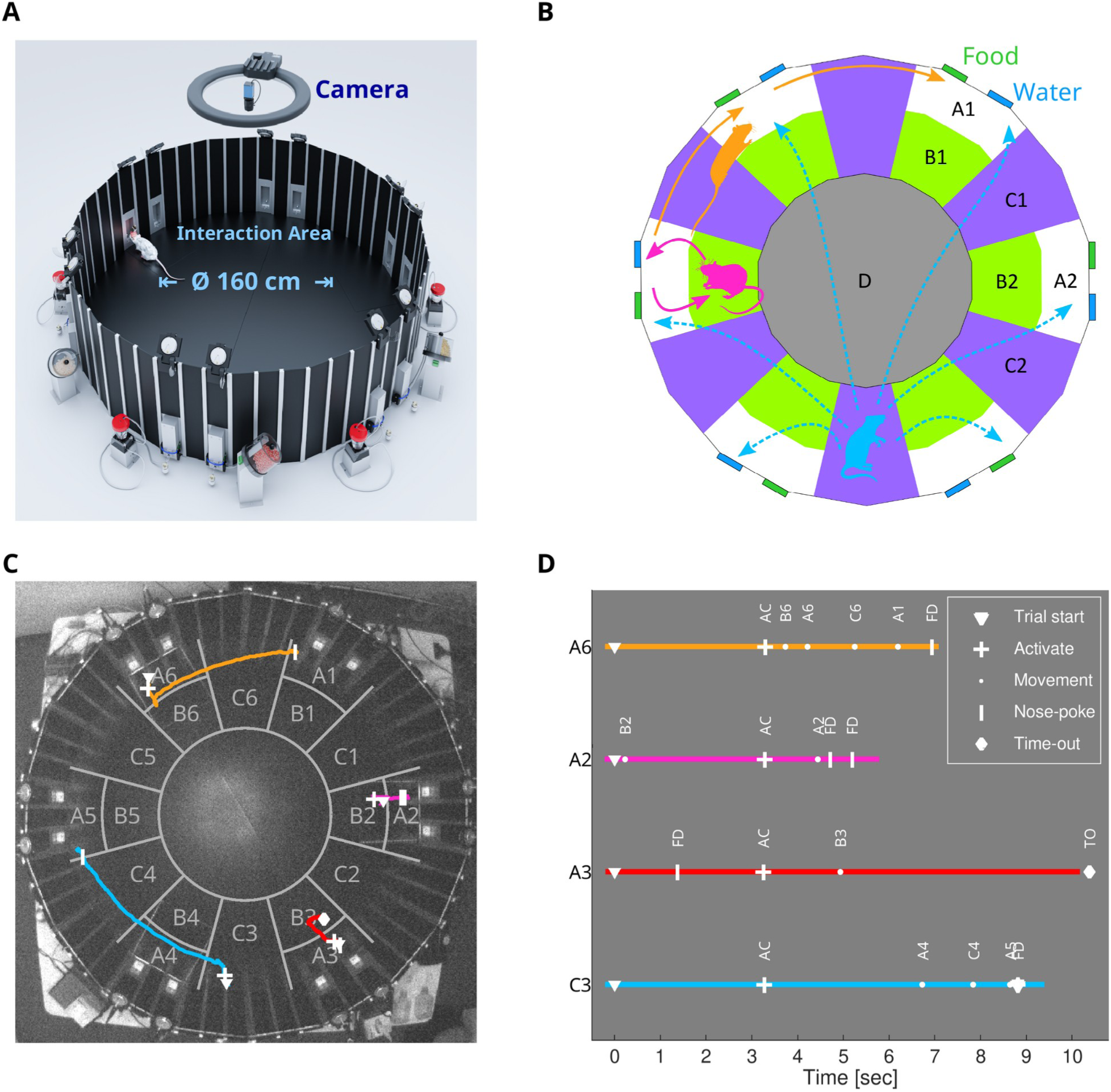
A foraging task formalized as a Markov Decision Process (MDP) (A) Rats were trained in the RIFF, an interactive arena that can implement arbitrary tasks. The RIFF has six Interaction Areas (IAs). Each IA consists of two poking-ports for food/water delivery, and two loudspeakers to communicate information about the state of the environment to the rats. (B) Schematic of the task in the RIFF. This task consisted of self-initiated trials. After an attention sound, rat position was determined and assigned to one out of 19 specific sectors of the RIFF (marked in white, green, purple, and grey; called A, B, C, and D sectors below). After three seconds, a target Interaction Area (IA) was selected. The selection of the target area depended on the sector in which the rat was found as well as on the previous port from which the rat collected reward. The rules by which the target area was selected were designed to make the rat move all the time. If the rat was in an A sector, the next area was the one in which the rat was positioned unless it got reward from that area in the previous trial, in which case the target area was the clockwise neighbor (yellow). If the rat was in a B sector, the next area was the near A sector (red). If the rat was in a C sector, the next target was selected randomly among the six Interaction Areas (cyan). The rat had 20 seconds to reach this IA and nose-poke there to get the food or water reward. The target location was communicated via target-specific sounds. Nose-pokes elicited rewards (when correct) and feedback sounds (one sound for correct nose-pokes, and another for incorrect nose-pokes). Following feedback, a new trial was elicited unless the rat positioned itself in sector D. With 10% probability, a warning trial was initiated instead of a rewarded trial, in which case any nose-poke elicited a mild airpuff. (C) Four trials from one session. In this session, the rat used all available reward strategies (shown in orange, magenta, and light blue, colors corresponding to the schematic (B); a warning trial is displayed in red). Each trial begins with an attention sound (corresponding rat location indicated by ▼). After 3 seconds, the target interaction area (resp. warning trial) is selected and a target (resp. warning) sound is played (“activate” action; location indicated by +). The rat then has 20 seconds to move to the target IA to collect food or water (resp. has to avoid nose-pokes for 5 seconds in a warning trial; nose-pokes indicated by ❙; time-out indicated by ♦). Note that this rat had learned to barely move in the 3 seconds between trial start (▾) and target selection (+). (D) Time course of the four trials from (C). The sector in which the rat was placed during trial start is denoted at the left end of each trial. Trial start is indicated by ▼; movement actions by ● with their target location such as A6, B6; the “activate” action by + and AC; nose-pokes in a port by ❙ and FD, and time-out by ♦ and TO.

We start the analysis on the first day in which the rats were exposed to the full task, after a familiarization period during which rats were exposed to simpler tasks in the arena. Rats quickly learned the task (see Jankowski et al. (2023) for an analysis of the first two days of task performance). Over the first ten days of training, the percentage of successful trials out of all trials more than doubled, from 35% on the first day (average across 5 rats; range 27% to 44%) to 77% on the tenth day (range, 72% to 83%). We used a linear mixed effects model to model success rate as a function of day of training with random intercepts and random slopes for rats. This model showed a significant main effect for “day of training” (estimate, 0.033 ± 0.006; t(32) = 5.46; p = 5.2 × 10^-6^). Thus, on average, success rates increased by 3.3% percentage points from one training session to the next over the first ten days.

### Rat behavior as actions in a Markov Decision Process

In order to model rat behavior at a fine-grained level, we modeled the task as a Markov Decision Process. Markov Decision Processes (MDPs) are widely used in reinforcement learning (Sutton and Barto 2018). An MDP describes the environment as a finite but potentially large set of states, in which an agent – in our case, a rat – performs actions.

An action taken by the animal may transition the environment into a new state, and these transitions are potentially associated with the emission of a reward. Importantly, these transitions are “Markovian” – they depend only on the current state and on the current action selected by the animal, and not on previous states and/or actions. In the task used here, the RIFF was discretized into 19 sectors (Fig. 1B; for details see Methods). Normally, IAs were inactive and did not provide food or water upon nose-pokes. However, 3 seconds after trial initiation, the position of the rat determined one IA that was activated. A subsequent nose-poke in the activated IA was rewarded (provided it occurred before a 20 seconds timeout). The rule by which IAs were selected for activation depended on the sector occupied by the rat at trial initiation. Thus, trials are named as A, B, or C trials. The identity of the activated IA was communicated to the rats via sounds that were specific to each IA, presented from the loudspeakers of the activated IA.

In addition to these trials, at the time of trial initiation, a “warning” trial could be selected with a probability of 0.1. During such trials, any nose-poke elicited a mild airpuff. Warning trials ended after 5 seconds without nose pokes, or following a (incorrect) nose-poke.

For our MDP, the states were defined by the current position of the rat, the currently active IA, the previously active IA, and the current trial type (see Methods). We modeled rat actions in the MDP as 19 “movement” actions that moved the rat into the 19 sectors of the RIFF; nose-poke actions for the water and food port at each IA, that led to a state with no active IA; “activate” action that occurred at the time in which the RIFF logic selected the active IA; and a “timeout” action that occurred when the rat avoided poking for a long time (20 s in a regular trial, 5 s in a warning trial) and led to a state with no active IA. Reward was given whenever a nose-poke occurred in an active IA. Overall, the model consisted of 3724 states and 23 actions. However, only between four and twelve actions were available in each state (see Methods). Note that, strictly speaking, the “activate” and “timeout” actions occur due to the passing of specific time intervals, not due to an explicit rat action. However, we conceptualize the fact that the rat stays in its position in order to activate a trial (resp. to wait for a trial’s time-out) as an action. Some state-transition probabilities and the respective rewards are illustrated in Table 1.

**Table 1.** A partial state transition table for three states. Several actions are available at each state. State names indicate the rat position, currently active interaction area (IA), previously active IA, and currently active trial type. A reward is available at an active IA. For example, state (B2, 0, 3, A) means that the rat is in sector B2, no IA is active, interaction area 3 was used before, and the last trial type was A. Likewise, (A2, 2, 3, A) means that the rat is in sector A2, interaction area 2 is active, interaction area 3 was active before, and the current trial type is A; (A2, 2, 3, P) means that the rat is in sector A2, and the current trial is a warning trial, so that an airpuff will be delivered upon the next nose-poke. Actions include movement actions to each sector of the arena (e.g., A2, C2, etc.; these are available only for neighboring sectors), trial activation and timeout actions (AC, TO), and nose-poking (FD; this is available only in “A” sectors). This MDP has 3724 states and 23 actions. The state and the action jointly determine a probability distribution on the next state P(S_out_ | S_in_, a). For example, the trial activation (AC) in state (B2, 0, 3, A) leads to state (B2, 2, 3, B) with probability 0.9 and to state (B2, 2, 3, P) with probability 0.1. A state transition due to an action is associated with a reward R(S_out_, S_in_, a). For example, poking for food (FD) in state (A2, 2, 3, A) leads to a positive reward of size 30. See Methods for more details.

| State in | Action | State out | Probability | Reward |
| --- | --- | --- | --- | --- |
| *(position, IA-curr, IA-prev, trial-type)* |  | *(position, IA-curr, IA-prev, trial-type)* |  |  |
| (B2, 0, 3, A) | A2 | (A2 0, 2, A) | 1 | -1 |
| (B2, 0, 3, A) | C2 | (C2, 0, 2, A) | 1 | -1.5 |
| (B2, 0, 3, A) | C3 | (C3, 0, 2, A) | 1 | -1.5 |
| (B2, 0, 3, A) | AC | (B2, 2, 3, B) | 0.9 | 0 |
| (B2, 0, 3, A) | AC | (B2, 2, 3, P) | 0.1 | 0 |
| (A2, 2, 3, A) | FD | (A2, 0, 2, A) | 1 | 30 |
| (A2, 2, 3, A) | TO | (A2, 0, 2, A) | 1 | 0 |
| (A2, 2, 3, P) | FD | (A2, 0, 2, P) | 1 | -30 |
| (A2, 2, 3, P) | TO | (A2, 0, 2, P) | 1 | 0 |

Observed rat trajectories could be decoded into a sequence of MDP states and MDP actions. Four examples of individual trials performed by one rat in the same session are displayed in Figs. 1C and D. Each trial starts with an attention sound, indicated by ▼. After 3 seconds, the target interaction area (resp. warning trial) is selected and a target (resp. warning) sound is played; this is the “activate” action, indicated by +.

An example of an “A” trial is shown in Figs. 1C and 1D in orange. The rat was located in sector A6 at trial start and stayed there until target selection (interpreted as “activate” action). According to the rules governing target selection, the selected target IA was in sector A1. The rat then moved to B6, back to A6, and then via C6 to A1, where it poked into the food port and received a reward. A “B” trial is shown in magenta in Figs. 1C and 1D. After trial start, the rat moved to sector B2. This resulted in the activation of the IA in sector A2. The rat then moved to A2, poked and received a reward. A “warning” trial is shown in red. during this trial, the rat correctly refrained from poking into any port (time-out indicated by ♦). Note that the rat nose-poked without effect about one second after trial initiation, before the trial type was determined. In the last example (a “C” trial; cyan), the “activate” action occurred with the rat in sector C3, leading to the activation of the IA in sector A5. The rat moved through sectors A4 and C4 to A5, poked and received a reward.

### Policies under information constraints

A policy is a rule prescribing the probability of taking an action in every given state. MDPs have optimal policies – rules that reach the maximal reward rate possible. However, animal behavior often deviates from the optimal policies. Admittedly, we do not even know whether rat behavior is controlled by a policy, let alone an optimal one – action selection might depend on more than just the current state as we define it. However, as shown below, policies may approximate well observed rat behavior, and therefore for the rest of the paper, we will describe rat behavior using policies.

It is a ubiquitous observation that animals take many actions that are not optimal. Indeed, we found that rats often selected different actions when reaching the same state at different times during a session. Action distributions for some states are shown in Supp. Fig. 1. To describe such behavior, we used policies that are stochastic: in each state, there are many candidate actions that have a positive probability.

In complex tasks such as ours, directly estimating policies from observations is essentially impossible, because this requires estimating an exceedingly large number of probabilities from observations. While only 5 or 6 actions are available in most states, a policy still comprises 18788 potentially non-zero probabilities that need to be estimated from the data. However, a typical recording session consists of only 1500 state-action pairs. We therefore use an indirect approach for estimating policies. We compute a rich, parameterized pool of policies and select the one that best describes rat behavior.

The policies that we consider optimize a trade-off between *value* and *informational cost*. The *Value* of a policy *π* is simply the expected future reward, exponentially discounted as is the standard usage in reinforcement learning. The *informational cost* (IC) of a policy is a less standard quantifier. Formally, it is the expected future deviation of actions selected according to *π* from a default, task-agnostic policy. The task-agnostic policy was a uniformly random policy over all accessible actions in each state, but it may be selected as any policy in which action selection does not depend on the state, or depends on the state in ways that do not involve the task.

To understand the meaning of the IC, note that it is low when actions are selected completely randomly in each state. It is high when action selection is biased towards one, or a few, specific actions in each state. A policy with a high IC may require a precise determination of the state of the rat, as well as storing in memory the details of the exact action to select in each state. It is in this sense that the informational cost of such policies is high.

These two properties of policies – Value and IC – place every policy on an IC-Value plane (Fig. 2C). It is easy to conceive of bad policies (low reward) with high IC, but there may also be good policies with small IC. it turns out that for a given IC, there is a finite maximal Value that can be attained. This maximal Value is clearly a non-decreasing function of the IC that can be shown to be convex (Tishby and Polani 2011). Thus, the IC-Value plane is divided into two regions by a monotonically increasing, convex boundary curve that denotes, for each IC, the largest possible Value that policies with this IC can have. The boundary itself is achievable, and policies falling on the boundary are *information-limited optimal policies*: they have the largest possible Value for their IC. As will be shown below, these policies describe rat behavior well with a small number of parameters.

**Fig. 2.**
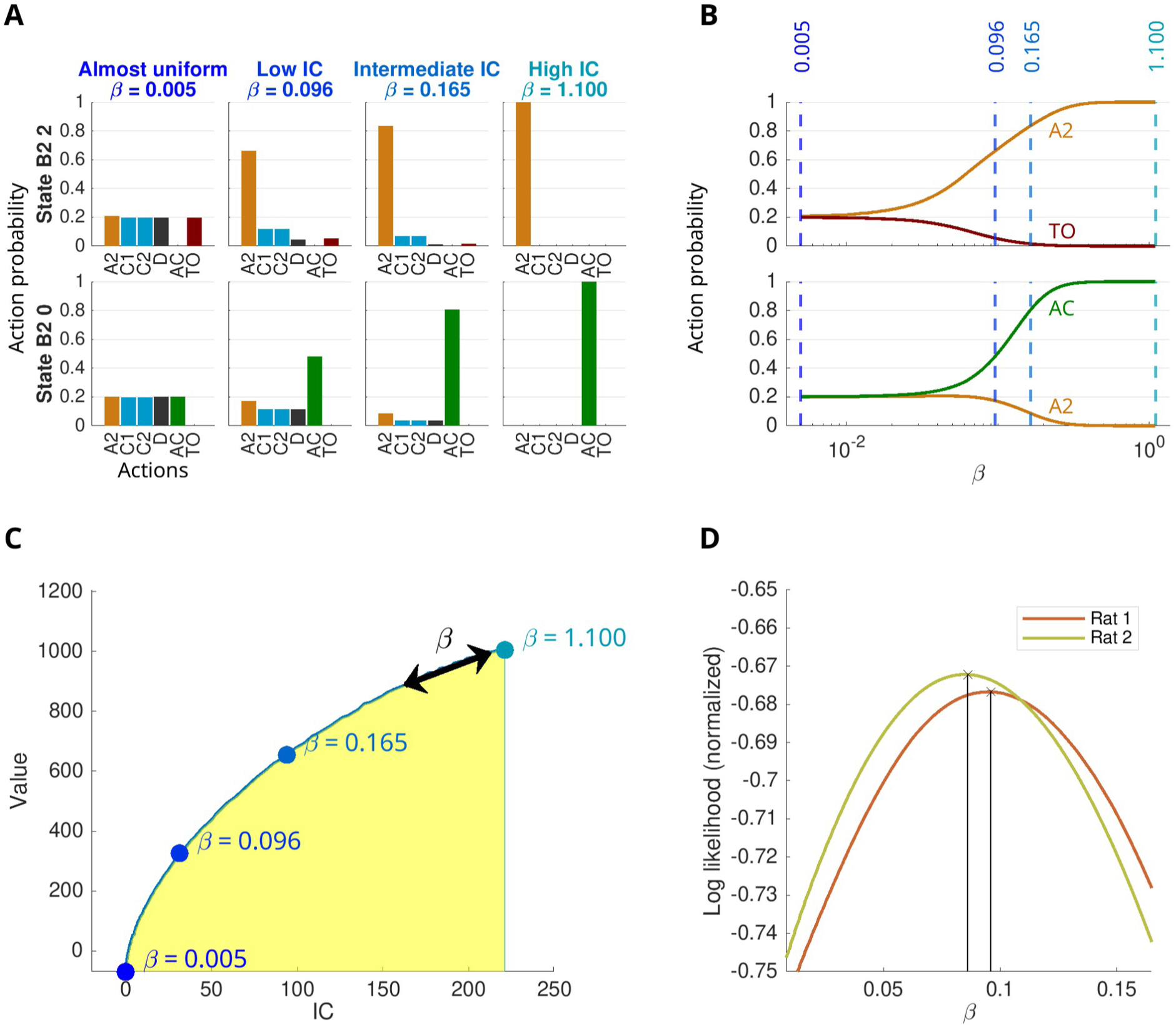
G learning creates MDP policies with prescribed informational complexity. (A) Examples of some state-dependent action probabilities of information-limited optimal policies. The probabilities of six actions are shown in two states, from information-limited optimal policies at four values of *β*. (Top) In the state abbreviated ‘B2 2’, the rat is in area B2 and reward is available when poking at the A2 interaction area (other state components omitted for clarity). The best action is ‘A2’ (i.e., go to A2), after which poking for the reward will be possible. (Bottom) In state abbreviated ‘B2 0’, the rat is in area B2 and no reward is available. Therefore, the best action is ‘AC’, i.e., activate. This will move the MDP into state B2 2. The four policies correspond to a very low *β*, resulting in an almost uniform policy; two intermediate values of *β* in which the probabilities of the optimal actions increase but are not yet 1; and a high *β*, resulting essentially in an optimal policy with probability zero for the non-optimal actions shown here. (B) The probability of each available action is a function of the parameter *β*, that controls the trade-off between information costs and MDP costs. The dashed lines indicate the policies displayed in (A). (C) Each policy defines a point in the IC-Value plane. Optimal policies under information constraints realize the maximum value at their IC, and therefore only IC and Value combinations in the yellow area are possible. The blue dots indicate the policies shown in (A) and (B). (D) Behavioral sessions are translated into a sequence of states and actions (compare Fig. 1D). The two curves show the log-likelihoods per action, for two sessions from two different rats, computed for the information-limited optimal policies at every value of *β*. For each policy, the log-likelihood per action is the average of the log probabilities for the observed actions, as specified by the policy with that *β*. Each session is then characterized by the *β* that maximizes the log likelihood.

The task of finding information-limited optimal policies can be formalized as the optimization problem of finding policies *π* that minimize the “free energy” *F^π^*

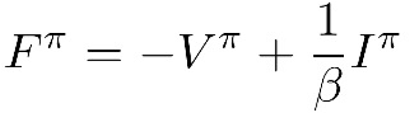

where *V^π^* is the Value of the policy, *I^π^* its informational complexity, and *β* is a trade-off parameter that balances the relative importance of the policy’s Value and IC (see Methods for details). A given *β* implicitly determines a constraint on IC such that the resulting Value is maximal among all policies that fulfill the constraint. When *β* = 0, the implicit constraint on the IC requires it to be 0. In consequence, the policy that minimizes the free energy is the default policy, the animal behaves randomly, and reward is minimal. Larger *β* results in a relaxation of the constraint on the IC, and the policies that minimize the free energy have higher Value. For large enough *β*, the implicit constraint on the IC is weak enough so that it is fulfilled by optimal policies of the MDP, and the policy that minimizes the free energy converges therefore to an optimal policy. We used a modified version of G learning (Fox, Pakman, and Tishby 2016) to create a set of information-limited optimal policies for a range of *β* values.

Examples of some action probabilities in information-limited optimal policies for four different values of *β* are shown in Fig. 2A. At low values of *β*, all accessible actions have very similar probabilities (i.e., the policy is very close to uniformly random action selection). As *β* increases, the probability of the optimal actions in each state increases, and converges to 1 at very high *β* values. Action probabilities for two actions in two states as a function of *β* are shown in Fig 2B, suggesting that the exact dependence of action probabilities on *β* is non-trivial. The optimal information-limited policies created by G learning fall on the boundary curve in the information-value plane (Fig. 2C; the position of the four example policies shown in Fig. 2A are marked).

We encoded each session performed by the rats as a sequence of MDP states and actions (called ‘the behavioral trajectory’ for that session). Since a policy is a table of probabilities of actions in each state, we computed the log likelihood of a trajectory given a policy as the sum of the log probabilities for the observed actions along the trajectory (see Methods). For the family of information-limited policies parametrized by the trade-off parameter *β*, the likelihood is a function of *β* (see Fig. 2D for examples of likelihood functions). For each behavioral session we found the best-fitting policy among all optimal information-limited policies. We identify these policies by their tradeoff parameter *β*.

When the best-fitting *β* is close to zero, the complexity is close to zero and rat behavior is best described by a policy in which actions do not depend much on state, suggesting that the rat is not performing the task. When the tradeoff parameter *β* is higher, complexity is higher, the policy is different from the default policy, and we further know that this policy has optimal Value among all policies with that IC. Such a policy requires the rat to identify the different states and to select reasonably its actions, suggesting it is under task control. We therefore operationally define the *β* value of the best fitting policy as the **“task alignment”** (TA) of the rat.

### Prolonged increases in task alignment during training

We used day-by-day changes in task alignment to study learning. We first analyzed task alignment in the first month of training (Fig. 3A). There was a significant main effect of day of training (linear mixed-effects model with random intercepts and slopes for rats; F(1, 70) = 93.5; p = 1.6 × 10^-14^). In fact, day of training and task alignment were positively and significantly correlated in each rat (ϱ > 0.56 in each rat, p < 0.027 in each rat).

**Fig. 3.**
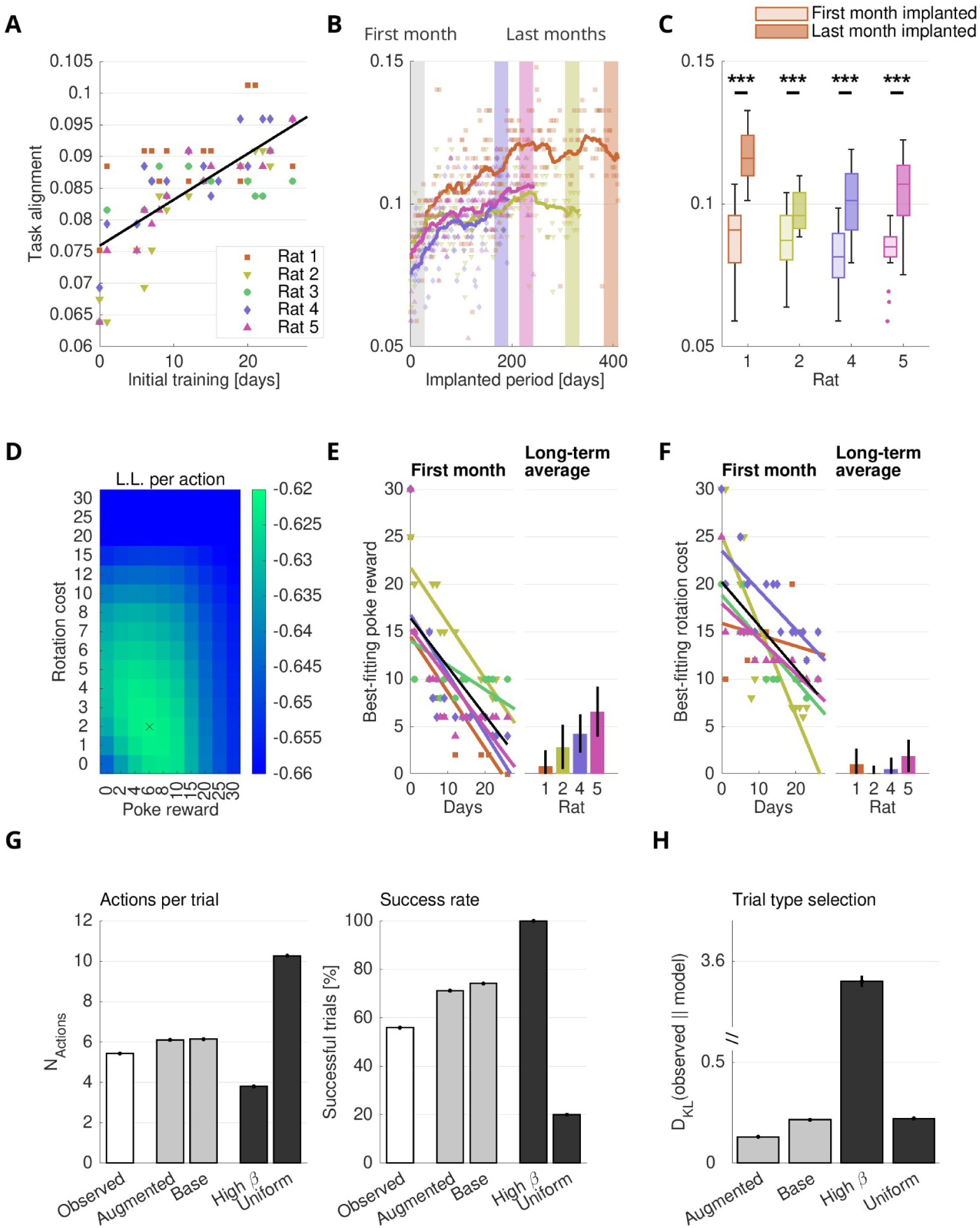
Information-limited optimal policies quantify learning and fit well behavior. (A) Task alignment increased over the initial 28 days of training in all five rats. Shown is the estimated *β* (i.e., task alignment) of rats for each session. (B) Task alignment continued to increase also after electrode implantation. Shown is the estimated *β* of four rats, for the entire period after electrode implantation. Dots depict individual sessions, continuous lines depict 28-day moving averages. Background patches mark the first 28 days after implantation (gray), and the last 28 days of the experiment in each of the four rats (color). (C) The task alignment was significantly higher during the last month of the experiment compared to the first month after implantation, in each rat. The box plots are based on the task alignment values of the individual sessions from (B). (D) Average log-likelihood was used to estimate the best-fitting parameters for the MDP augmented by poke reward and body rotation costs. (E) Estimated poke rewards decreased quickly during the first month of training. Left panel: Poke rewards for each session during the first month of training. Right panel: long-term averages of the poke rewards for the entire time period after implantation. Error bars indicate standard deviation. (F) Estimated orientation-costs decreased during the first month of training. Left and right panel as in (E). (G) Macroscopic statistics of simulated agents under best-fitting Information-limited optimal policies resemble observed behavior. The left panel displays the average number of actions needed to complete a trial over all sessions, compared with that determined for four policies: the default policy, an optimal policy, the best-fitting information-limited optimal policy, and the best-fitting information-limited policy for the augmented model with poke rewards and body rotation costs. The last two policies were estimated for each session separately. Error bars show s.e.m. Right panel, same as left, but for success rates. (H) The D_KL_ between the trial type distribution (how many A, B and C trials occurred in each session) and that determined from the simulated trajectories of the same four models as in (G).

This long-lasting learning effect could simply reflect an increase in success rates over time. However, success rates showed a markedly shallower and more variable increase as a function of day of training (Supp. Fig. 2A). Modeling jointly standardized task alignment and standardized success rate as a function of day of training (linear mixed effects model, fixed effects: day of training and observation type with interaction; random intercepts and slopes for rats) revealed a significant main effect of day, as expected (F(1, 140) = 9.20; p = 0.0029) as well as of observation type (i.e., “task alignment” vs. “success rate”; F(1, 140) = 4.62; p = 0.033). Importantly, the slope of task alignment as a function of day was significantly larger than that of success rate (significant interaction of “day” and “observation type”; F(1, 140) = 6.51; p = 0.012). We conclude that task alignment described the learning process of the rat in a finer way than success rate, presumably because task alignment takes into account all rat actions at a high temporal and spatial resolution.

Following a lengthy training (about 4 months), rats were implanted with electrodes for extracellular recordings, disrupting for a while their task performance (see Methods). After implantation, four of the five rats performed the task for many months (193 days, 242 days, 333 days, and 410 days, respectively). Task alignment for the four rats over the full period is displayed in Fig. 3B. We observed a clear increase of task alignment up to approximately 200 days after implantation in each individual rat. In the two rats with the longest observation periods, task alignment plateaued after 200 days. Task alignment consistently increased during the post-implantation period (Fig. 3C; task alignment in the first 28 days after implantation compared to the last 28 days of testing: two-sample t-test, P < 10^-8^ in each rat).

While success rates after implantation continued to increase for a few weeks in each rat, they reached a plateau earlier than task alignment, after about 50 days (Supp. Fig. 2B), while task alignment continued to increase for at least an additional 100 days. To capture the dynamics of success rates compared to task alignment over these long time periods, we z-scored both task alignment and success rate, and then modeled the standardized measurements in three time windows: the entire implanted period, the time period after day 50, and the time period after day 150 (linear mixed effects models with random intercepts and random slopes for rats). In all time windows, the estimated slope of task alignment as a function of day was positive and significant (all p < 0.0012; see Supp. Table 1), showing that task alignment continued to increase even after day 150. The estimated slope of success rate as a function of day was significantly smaller in all three time periods (significant interaction between day of training and observation type in each time period, all p < 4 × 10^-8^; see Supp. Table 1). Over the entire implanted period, the slope in success rate was positive, but about 3 times smaller than the slope in task alignment (2.30 × 10^-3^ standardized units/day vs. 6.42 × 10^-3^ standardized units/day). In the later testing periods, after day 50 and after day 150, the estimated slope in success rate was in fact negative, while task alignment continued to increase in both cases.

Thus, over these very extended periods of testing, task alignment revealed a continuing learning process that was not apparent when considering only success rate.

### Estimating internal costs and rewards

Two aspects in the behavior of the rats changed over the long training period. First, at the beginning of training, rats tended to nose-poke in the absence of actual rewards, when no IA was active. This was likely due to the way their behavior was shaped before they were exposed to the task (see Methods). Second, as time progressed, rats tended to activate IAs from B areas more often, which required them to rotate by 180 degrees (to move from the A to the B area, and then to return to the same A area; this was encouraged, see below and Methods).

To capture these changes, we extended the model by refining the MDP and by adding two parameters to the reward function. First, we added internal costs for body rotations. To do that, we introduced a body orientation to the model (discretized to four directions) so that each state of the original model gave rise to four states in the new model, each with a different body rotation of the rat. We then added a small cost for transitions that resulted in changes of the body orientation. Second, we modeled the tendency of the rat to poke even when no reward was provided by adding small rewards for any poke, even when the poked IA was not activated so that the poke did not result in a reward. These small rewards can be interpreted as ‘internal rewards’ that drive an action which by itself doesn’t result in an externally-delivered reward. This model has a total of 14896 states (the previous 3724 states times four orientations; see Methods). When the costs for body rotation and the reward for a nose-poke are both set to 0, this model can be reduced to the previous one. We refer to this MDP as the augmented model.

We considered the poke reward and body rotation cost as two additional parameters of the model, together with the tradeoff parameter *β*. We computed the information-limited optimal policies at all *β* for each combination of 10 nose-poke rewards (0–30) and 15 body rotation costs (0–30; see Methods). As before, we determined the policy that had the maximum likelihood among all these candidates, maximizing over the three parameters (*β*, body rotation cost, and nose-poke reward). An example set of log-likelihoods as a function of poke reward and body rotation cost, at a fixed *β*, is shown in Fig. 3D.

In all rats, the best-fitting poke reward sharply decreased during the initial month of training (Fig. 3E), corresponding to the decreased propensity of the rats to perform useless pokes. There was a significant effect of day of training (linear mixed effects model with random intercepts and slopes for rats; F(1, 70) = 58.3; p = 8.6 ×10^-11^), with the best-fitting poke reward declining on average by about 0.5 units/day. For the period following implantation, the best-fitting poke reward was much more stable (same model; no significant effect of day of training (F(1, 637) = 0.0051, p = 0.94).

The best-fitting rotation cost also sharply declined over the initial month of training (Fig. 3F), reflecting the increase in sharp body turns made by the rats. The best-fitting rotation cost decreased on average by 0.46 units/day (significant effect for Day; linear mixed effects model with random intercept and random slope for rats; F(1, 70) = 14.6; p = 2.8 ×10-4). Since the rats still used the B areas only rarely after the initial month of training we actively encouraged rats to use them (see Methods). In consequence, there was an additional decrease in body rotation costs between the end of the first month of training and the period following implantation, a few months later (range of differences, 6.93 to 14.5 units across four rats; all P < 10^-90^), indicating that rats further learned to make sharp body turns later during training.

### Information-limited policies approximate observed rat behavior

In the previous sections, changes in the best-fitting policies were used as evidence for changes in behavior (and therefore learning). However, the policies that we match to rat behavior are only a small subset of all possible policies, and are unlikely to be the precise policies followed by the rats. We want to study the resulting bias -how similar are these policies to the observed rat behavior?

MDP models are generative: a given policy can be used to compute simulated trajectories of agents following the policy. Relevant averages of behavioral parameters from the simulated trajectories can then be used to compare agents with observed data. We used this approach to compare the observed rat behavior in each session to four different policies: The best information-limited policy for the base MDP, fitted to each session (with a single parameter, *β*; as in Fig. 2); the best information-limited policy for the augmented MDP, fitted to each session (with three parameters: *β*, poke reward, and body rotation cost; as in Fig. 3); and two control policies: the default uniform policy (providing a lower bound on performance) and the optimal policy derived from the base model (providing an upper bound on performance; see Methods). We examined three statistics of behavior: the number of MDP actions needed to complete a trial (Fig. 3G, left), the fraction of successful trials (Fig. 3G, right), and the distribution of trial types (.e., the fraction of A, B, and C trials in each session; Fig. 3H). We compared the experimental distributions of these statistics with those resulting from the policies fitted to each session (See Methods).

The information-limited policies (both base and augmented MDPs) had consistently better fits to the observed data than either the default or the optimal policies (Fig. 3G and H; underlying data in Supp. Table 2; statistical comparison in Table 2). For example, rats needed on average 5.43 ± 0.046 actions per trial, while MDP trajectories in the best-fitting policy for the augmented MDP had on average 6.10 ± 0.023 actions per trial. The other two policies were farther from the observed data: for the default random policy, MDP trajectories had 10.27 ± 0.01 actions per trial, while for the optimal policy, MDP trajectories had 3.8 actions/ trial with very little variability (3.7977 ± 0.0005 actions per trial; see Supp. Table 2 and Table 2; significant main effect of type of model on actions per trial; F(4, 3030) = 8923; p < 10^-100^).

Likewise, success rates of rats were most closely reproduced by trajectories estimated from the best policies for the augmented MDP (see Supp. Table 2 and Table 2; significant main effect of type of model on success rate; ANOVA F(4, 3030) = 16359, P < 10^-100^).

**Table 2.** The 3-parameter model most closely resembles the observed data. Statistical comparison of “actions per trial” and “success rates” for four different models. The t and P values refer to paired t-tests.

| Actions per trial |  |  |  |  |
| --- | --- | --- | --- | --- |
| Model 1 | Model 2 | d' | t(606) | P |
| Observed | Augmented | -0.59 | -19.8 | 1.8E-67 |
| Observed | Base | -0.62 | -19.6 | 8.3E-67 |
| Observed | High beta | 1.4 | 35.4 | 1.0E-149 |
| Observed | Uniform | -4.3 | -102 | 0.00E+00 |
| Success rates |  |  |  |  |
| Model 1 | Model 2 | d' | t(606) | P |
| Observed | Augmented | -1.9 | -41.8 | 8.6E-181 |
| Observed | Base | -2.3 | -52.7 | 1.5E-228 |
| Observed | High $\beta$ | -5.5 | -135 | 0.0E+00 |
| Observed | Uniform | 4.5 | 108 | 0.0E+00 |

While statistics estimated from the trajectories of the best-fitting policies of the base MDP were very similar to those derived from the best-fitting policies of the augmented MDP, the difference in the goodness of fit between the two was significant in both cases due to the large number of sessions and trials (statistical analysis in Table 2; paired t-test for base vs. augmented MDPs; actions per trial, P = 5.8 x 10^-7^; success rate, P = 2.7 x 10^-87^).

We found similar results for the distribution of trial types: rats could freely choose among A, B, and C trials, independently in each trial (the distribution of trial types for rats and each model is listed in Supp. Table 3). The trajectories determined by the best fitting policies for the augmented MDP produced trial type distributions that most closely resembled those observed in rat behavior (Similarity of the observed trial type distribution and the distributions derived from the best-fitting policies quantified as the DKL between them; see Methods and Table 3; a significant main effect of type of model on the similarity of the observed distributions to the distributions from the simulated trajectories, F(3, 2424) = 11558, P < 10^-100^), with the policies of the augmented MDP outperforming all others (all P < 4 x 100^-60^; see Table 4).

**Table 3.**
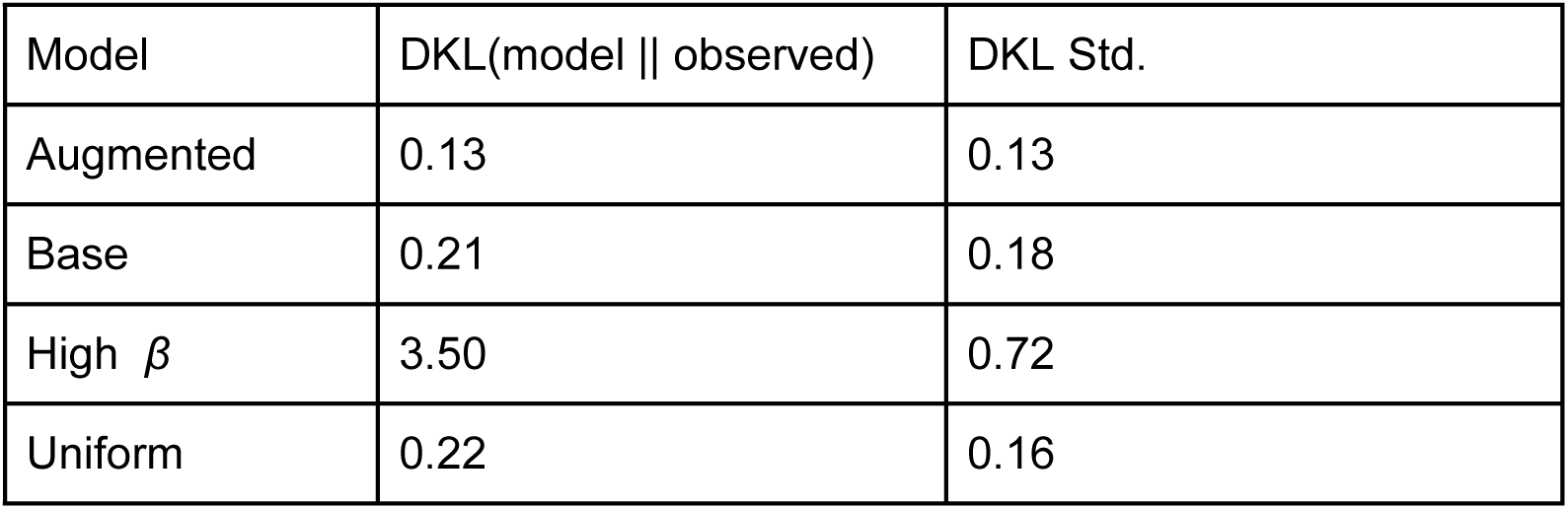
Comparison of trial type distributions between observed data and each model. DKL Mean/Std are D_KL_ of the model’s trial type distribution compared to the observed data.

**Table 4.** Comparison of the augmented model to the other models. The t and P values refer to paired t-tests between the models.

| Model 1 | Model 2 | T(606) | P |
| --- | --- | --- | --- |
| Augmented | Base | -18.3 | 3.92E-60 |
| Augmented | High $\beta$ | -127.0 | 0.00E+00 |
| Augmented | Uniform | -18.6 | 1.73E-61 |

### Insular cortex neurons correlate with within-session changes in task alignment

We wanted to study how task alignment changes within a session. We found that 10-minute sections were long enough for estimating stably the best fitting policy for the base model (without poke rewards and body orientation costs; this was justified by the fact that for the period following implantation, these rewards and costs were minimal).

We subdivided each session into ten-minute windows with 9 minute overlap between successive windows. We therefore estimated the task alignment, quantified as the *β* of the best-fitting policy, from the actions within each of these windows. The time course of task alignment within a session, averaged over all sessions of all rats, is shown in Fig. 4A. Task alignment increased on average during the first few minutes of a session, presumably while the rat calmed down after its introduction into the RIFF, and then held largely stable until the middle of the session. The second half of the behavioral sessions was more variable both within and between rats. While rats were not fully sated at the end of the behavioral sessions (they always consumed more food in their home cages following the behavioral sessions), they may have been less hungry by that time, and therefore behaved less consistently.

**Fig. 4.**
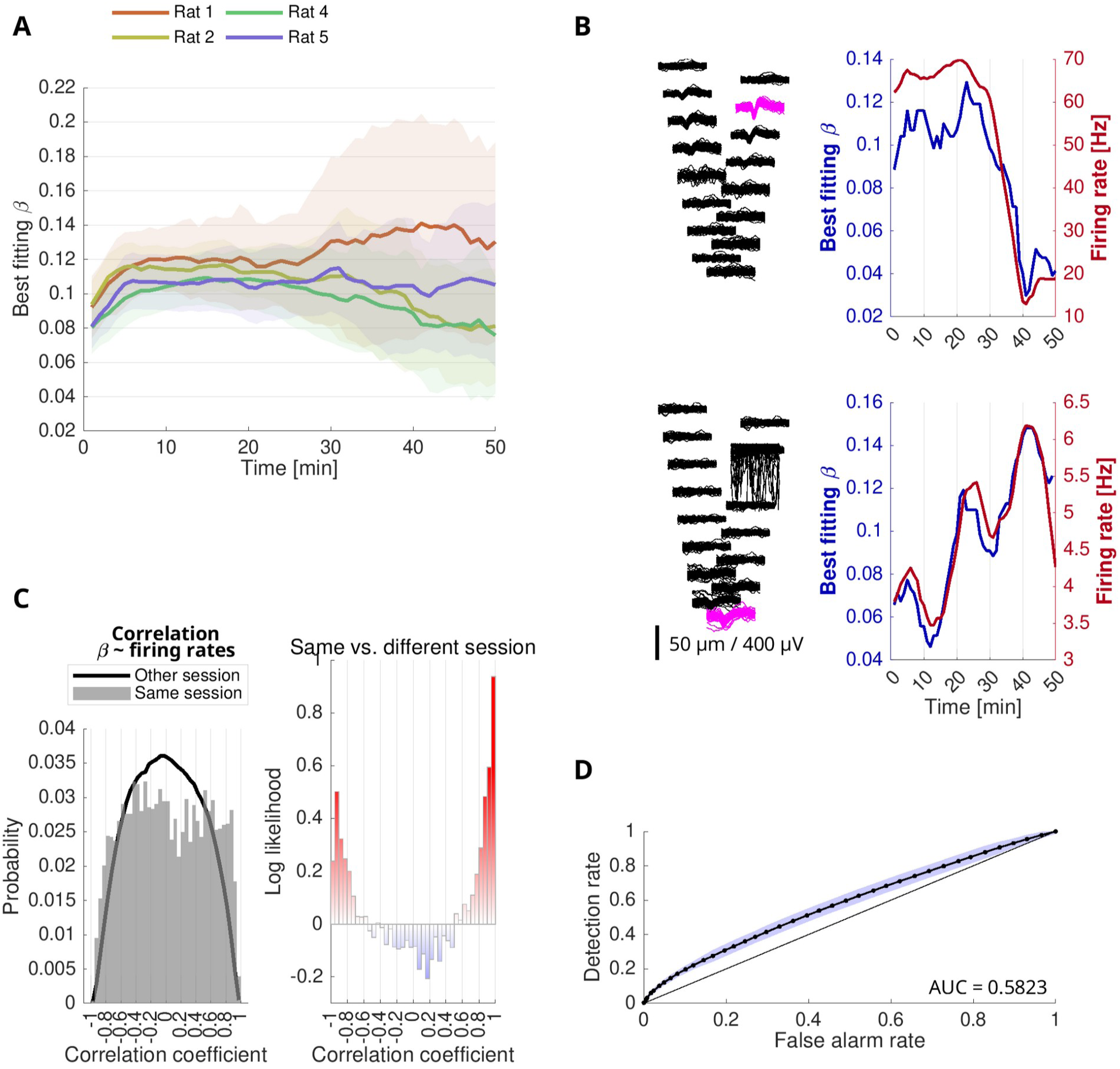
Insular cortex encodes task alignment. (A) Behavioral complexity within sessions estimated in overlapping 10-minute windows with a 1-minute shift between windows. Shown is the estimated *β* parameter for four rats, from all sessions during the implanted time period. The solid line indicates the mean, the shaded areas indicate standard deviations around the mean. (B) Examples of two neurons whose firing rates were highly correlated with *β* within a session. (Left panels) Spike waveforms as recorded on all 16 contacts of one side of the recording electrode shank. (Right panels) Average firing rates, and *β*, were estimated in overlapping 10-minute windows. (C) (Top left) Distribution of Pearson correlation coefficients between *β* and firing rates for all recorded neurons. (Bottom left) Control. distribution of correlation coefficients between firing rates of all units, and *β* from all recording sessions other than the session that the unit was recorded in. (Right) Comparison of distributions. Shown are log-likelihoods for correlation coefficients when comparing the distribution of correlations between *β* and firing rates from the same session, and other sessions. Correlation coefficients with high absolute value are more likely to originate from units and *β* recorded during the same session. (D) ROC analysis for correlation coefficients. Each point on the ROC curve corresponds to a probability threshold, and depicts fractions of correlation coefficients. The shaded area indicates the standard deviation around the ROC curve, estimated by a bootstrap procedure.

We compared the time course of task alignment in each individual session with firing rates of the neurons recorded in the insular cortex during the same session. The firing rates were computed over the same 10-minute windows used to compute task alignment (Fig. 4B; see Methods). The two examples in Fig. 4B show neurons whose firing rates had a high positive correlation with the fluctuations in task alignment in the same session. We calculated correlations between the firing rates of each neuron and the time course of task alignment in the session in which the neuron was recorded (Fig. 4C, top left; 305 sessions from four rats; 5576 units total). This distribution is compared with the distribution of correlations between the firing rate of each neuron and the task alignment in each of the other sessions (Fig. 4C, bottom left), showing that large positive as well as negative correlations between firing rates and same-session task alignment were over-represented in the data (Fig. 4C, right). An ROC curve analysis confirmed this finding (Fig. 4D), with an area under the curve (AUC) of 0.582 that was significantly different from 0.5 (bootstrap, 1000 samples, t(999) = 83.3, P < 10^-100^; see Methods). Thus, the firing rates of many neurons in insular cortex tracked the time course of task alignment during the behavioral tests.

High correlations between task alignment and neuronal firing rates could originate from units that responded to rewards, simply because time segments with higher task alignment would also have more rewarded actions. Indeed, repeating the same analysis using the correlations between the firing rates of each neuron and the average number of rewarded trials (computed in the same time windows) showed also an excess of high correlations, verified by an ROC analysis (Supp. Fig. 3A; AUC = 0.648; 1000 bootstrap samples, t(999) = 157; P < 10 ^-100^). Note that correlations between rewards and firing rates were predominantly positive, while for task alignment, both negatively and positively correlated neurons were over-represented.

We therefore analyzed the correlations between task alignment and firing rates, controlling for rewards. For that purpose, the firing rate of each neuron was modeled as a function of both rewarded trials and task alignment. We tested the additional information provided by task alignment beyond the contribution of the reward to the firing rate by comparing the goodness of fit (R^2^) of that model with the goodness of fit of models that used task alignment from other sessions (but the rewards from the same session; Supp. Fig. 3B; an exemplar unit is shown in the main plot). The surrogate goodness of fit tended to be lower than that of the true data (Fisher’s exact test for 95% upper ranks of R^2^, true vs. randomly permuted session session label, P = 6.6 × 10^-5^; two-sample Kolmogorov-Smirnov test for true ranks of R^2^ vs. mean of 100 randomly permuted session labels, D252,252 = 0.444, P = 1.5 × 10^-22^). Thus, for a significant fraction of insular cortex neurons, the task alignment explained firing rates beyond reward rates alone.

## Discussion

We introduce here a novel approach for a detailed analysis of rat behavior, based on Markov Decision Processes and notions from information theory. We derive, from first principles, a measure of task alignment that depends on behavior only and is easily determined from observed data. We show that task alignment provides a rich source of information about rat behavior over multiple time scales: rats refined their behavior progressively over more than one hundred days in a way that was not captured by standard measures of behavior, such as reward rates; on the other hand, we found that within-session minute-by-minute fluctuations in task alignment were tracked by firing rates of neurons in the insular cortex.

### Markov Decision Process models of complex behavior

In the MDP framework, rat behavior is encoded as a sequence of states and actions. There are many choices available for the level of details in the specification of the MDP. However, once the level of detail (largely, spatial and temporal resolutions) is selected, much of the data that defines the MDP is derived directly from the design of the task. The states we selected describe rat location (and orientation, in the augmented model) with some detail; the actions that are available to the rat in each state are easily defined; and the state transition graph is constrained by these two. These MDPs described rat behavior at a sub-trial level (Fig 3 G; average of 5.4 actions per trial, corresponding to 20.2 actions per minute), at the price of being somewhat large: some of the MDPs analyzed here had 14896 states and 23 actions. In future work, it may be possible to reduce the number of states by systematic state aggregation (Li, Walsh, and Littman 2006; Sutton and Barto 2018), or alternatively to improve the fit of the MDP to behavior by using a spatially more fine-grained model of the arena with fine-grained movement actions (Kazakov et al. 2022).

While states and actions are fully observable in our approach, the reward function is unobservable. We used simple forms of rewards - small negative rewards for any movement, and a large reward for food or water. We also introduced positive rewards for unreinforced pokes (to account for the large number of those early in the training) and negative rewards for body rotation (to account for the small number of “B” trials early in the training). Importantly, we are interested in the policies rather than in the reward sizes, and policies are quite robust to small changes in reward size. In fact, we turned this argument on its head: by using some rewards as free parameters, we could extract consistent and interpretable trends in the reward function (see Fig. 3).

Given an MDP, the main question that we address is that of identifying the rules by which the rat selects its actions. MDPs can be optimally exploited by using a policy, defined as a rule that selects the next action based on the current state only. The candidate policies have to be non-deterministic and to allow the selection of suboptimal actions, in order to capture actual rat behavior. Our goal was to find a policy that describes rat behavior well.

The direct estimation of state-action probabilities from observed behavior is essentially impossible because most state-action combinations are observed only infrequently. Instead, we use a pool of policies estimated as maximally rewarding with a constraint on their informational distance from a default, task-agnostic policy (with equal probability for all actions accessible from each state). This is accomplished using G learning (Fox, Pakman, and Tishby 2016), which minimizes a free-energy function to generate policies. In practice, for each MDP we consider in this paper, we computed information-limited optimal policies spanning the range between the default policy and the optimal policy. We then characterize the observed behavior by the policy with the maximum likelihood.

Informational constraints on policies have been used also in control theory, with the Kullback-Leibler divergence from a default policy serving to define the cost of control (Rubin, Shamir, and Tishby 2012). Moreover, in multi-task reinforcement learning, entropy regularization has been used to induce similarity between task-specific policies and a common, task-agnostic policy (Teh et al. 2017; Tirumala et al. 2022; Liu et al. 2021; also see Moskovitz et al. 2022). However, in these studies regularization using informational constraints is used to improve the speed and robustness of learning, or the overall performance and generalization of the agent. To the best of our knowledge, this is the first time that such policies are used as candidates for describing animal behavior.

Importantly, we do not claim that the rat computes any quantities related to the MDP, and we do not study how the rat learned these policies (if it did that at all). Indeed, both the neural and behavioral relevance of state-action values is debated (Miller, Botvinick, and Brody 2022; Bennett, Niv, and Langdon 2021; Roesch, Taylor, and Schoenbaum 2006). Instead, our goal here is to characterize rat behavior with parameterized policies, regardless of how these policies are learned by the animals or how the rats select actions.

The use of these policies as approximants of rat behavior can be criticised. For one thing, we do not know that rats follow policies: it may be (and is actually likely) that the rats base their action selection on previous states and actions. The way by which rat behavior deviates from policies can be usefully studied. For example, it may be that rats select actions that also depend on their previous action (e.g. by repeating it). If such dependences are short enough in time, they can be accommodated at least partially into this framework by including past actions in the state. A model of this kind will have substantially more states, but could still be treated with the approach that we describe here.

Next, the origin of the suboptimal actions in our approach is assigned solely to suboptimal action selection (so to the decision making process). However, suboptimal actions could also arise from failure in identifying correctly the state (e.g. location in the arena). Models with uncertainty in the state belong to a different family, that of the partially-observed MDP (POMDP). POMDPs may be used to model constraints on sensation (Tishby and Polani 2011), by explicitly modeling partial observations (Spaan 2012). However, POMDPs are much more complex to work with, and they do not have a general optimality theory. We therefore preferred to work with the simpler MDPs theory.

Finally, it is also important to recognize that since we compare behavior with a limited set of policies, there may be (and likely there are) differences between the observed behavior and the best-fitting policy. Thus, we are certain to have a bias in the estimation of the best-fitting policy.

In spite of this bias, an important result of this paper consists of the demonstration that the information-constrained policies fit observed behavior significantly better than either of the task agnostic or optimal policies. The behavior of an agent following the MDP using the best-fitting policy is surprisingly close to observed behavior. Indeed, we show that many macroscopic observables that summarize rat behavior match well those of our selected best-fitting policies. The good fit between the best-fitting policies and observed behavior is evidence that our choices (using MDPs and policies to describe behavior, the reward functions we developed, and the use of informational constraints to parametrize the policies) do align with those of the rat. The information-constrained optimal policies form therefore a useful class that is rich enough to realistically describe the behavior of real actual rats performing a real complex task. In that sense, this approach allows us to peek into the rat mind.

### Task alignment, behavioral refinement, and learning

The informational constraint we use consists of the DKL between that policy and the default policy that assigns equal probability to all available actions. This default policy does not require the rat to identify its state in any details, so that action selection is trivial. In that sense, the default policy is simple. The optimal policy, on the other hand, requires the rat to know exactly in which state it is, as well as the ability to match the state against the policy in order to select the best action, imposing a substantial load on perception and memory. In that sense, the optimal policy is complex. We interpret the tradeoff between informational constraints and reward as a tradeoff between the resources required to implement the policy on the one hand and the expected reward on the other hand.

We defined task alignment using the tradeoff between informational constraints and reward that is encoded in the free energy function, using the *β* tradeoff parameter of the best fitting policy. Task alignment increased over the first month of training, implying that the best-fitting policy became more similar to the optimal policy, with finer distinctions between states and higher probabilities for useful actions in each state. This increase was more pronounced and less variable than the concurrent increase in trial success rates. Importantly, changes in task alignment reflect changes in behavior. The long-term increase in task alignment (up to 200 days after implantation) is evidence that additional learning was ongoing even in highly trained animals, resulting in behavioral refinement: rats learned to align their behavior more closely with the goals of the task, even if this produced only a mild effect on success rates.

Another way by which we documented learning over extended periods of time consisted of extending the MDP by introducing two internal cost parameters, poking reward and orientation cost. We then selected the best information-constrained policies in this augmented family, so that the best fitting policy now selected both the best fitting cost parameters and the *β* parameter. While the goodness of fit of the policies in this augmented family was only marginally better than that of the original MDP, the cost parameters showed a consistent trend across animals. Thus, with due caution, our approach may access information regarding internal costs and rewards governing rat behavior, and how they change throughout learning.

### Short-term task alignment is reflected in neural activity

Motivational states of animals are known to vary within experimental sessions. These changes likely reflect internal, unobserved changes in motivational states, and are sometimes described as changes in task engagement (Ashwood et al. 2022; Grohn et al. 2024; San-Galli et al. 2018). We therefore estimated task alignment in 10 minute time windows, using a few tens to a few hundreds of actions performed by the rats in these time windows. We consider the fluctuating values of task alignment computed this way as a way of quantitatively gauging the engagement of the rat in the task. When the rat is highly engaged, its behavior should approximate better that of an optimal policy, consistent with policies that have higher task alignment. Rats that are uninterested in the task are expected to show behavior consistent with low-IC policies, and to fail to collect much reward, consistent with policies that have lower task alignment.

We then used the short-term task alignment as a regressor for neuronal activity. Many neurons in the insular cortex were strongly correlated with task alignment. However, neurons in insular cortex show fast phasic responses to stimuli in various modalities, such as learned food cues, licking, food (Livneh et al. 2017) and sounds (Jankowski, Karayanni, et al. 2023). Remarkably, this correlation held for many neurons even after controlling for the rate of reward acquisition. Indeed, insular cortex has been linked to both low-level and integratory single- and multi-modal brain functions, including the gustatory system, action selection, and integration of motivational state (Vincis and Fontanini 2019; Livneh and Andermann 2021; Livneh et al. 2017). Activity fluctuations of insular neurons at slow time scales have been shown in the context of physiological states such as thirst (Livneh et al. 2020) and visceral malaise (Gehrlach et al. 2019). Here we suggest the existence of similar activity fluctuations over slow time scales that encode the internal variable of task engagement.

## Conclusions

We describe here a novel approach to quantify rat behavior that takes into account each and every animal action. We explicitly compute information-constrained optimal policies and show that this family of policies fit observed behavior surprisingly well. The notion of task alignment that we introduce provides a novel window on learning: task alignment continues to increase even when success rates stagnate, rigorously demonstrating behavioral refinement even without an increase in reward acquisition. Furthermore, we report correlations between insular cortex neurons and task alignment on a slow timescale, suggesting that neurons in insular cortex encode task engagement.

## Methods

### Rat arena and task

The behavioral arena (the RIFF) has been described previously (St+ task; Jankowski, Polterovich, et al. 2023). Briefly, the RIFF is a circular arena (diameter 160 cm) with six interaction areas (IAs) equally spaced along its circumference (Fig. 1A). Each IA consists of two ports and two loudspeakers. One port in each IA is equipped with a pump for liquid delivery, the other with a food pellet dispenser. Each port is equipped with an airpuff valve. The position of the rat is identified by a ceiling-mounted camera. The RIFF is controlled by a Matlab program (the code is publicly available at https://github.com/jniediek/RIFF_publication) and can implement tasks with arbitrary logic. We focus on tasks that have the structure of a Markov Decision Process (MDP; Sutton and Barto 2018). An MDP consists of a fixed number of well-defined states and actions together with a state transition matrix, which may lead to reward (positive and negative). Rat actions lead to state transitions and rewards are delivered depending on actions and state transitions.

For a full description of the St+ task, see Jankowski et al. (2023). In short, the arena was divided into 19 sectors A1–A6, B1–B6, C1-C6, and D (Fig. 1B). A trial was initiated by an Attention (ATT) event, marked by a sound. At an interval of 3 seconds after the ATT event, the upcoming target IA was determined based on the rat current and previous position, and the Target (TGT) event was triggered. At the TGT event, an IA-specific sound was presented at the target IA. The rat had to move to the target IA and nose-poke in one of the two ports at that IA. The Feedback (FDB) event was triggered by the first nose-poke into any port. If the nose-poke was at the correct target IA, food/water was provided and a “correct” sound was presented. If the nose-poke was not at the target location, no reward was provided and a “mistake” sound was presented. If the rat did not poke at any port within 20 seconds after the TGT event, a time-out occurred, and the “mistake” sound was presented. In a randomly selected fraction of all TGT events, instead of activating an IA, a “warning” trial was initiated and a “warning” sound was presented. In a warning trial, a nose-poke into any port resulted in a mild air-puff together with a specific sound. The timeout in a warning trial occurred 5 seconds after trial initiation, and a “safety” sound was presented to indicate the end of the warning trial. At an interval of 3 seconds following the FDB event, a new trial was initiated by an ATT event, unless the rat moved to the D area. Trials restarted once the rat moved out of the D area.

Rewards consisted of 45 mg food pellets with one flavor per IA (six flavors; TestDiet, Richmond, IN, USA), or flavored water (four flavors). The amount of food pellets provided as a reward depended on the area where the current trial was initiated: for trials triggered from an A sector, the reward was one food pellet/unit of liquid. In trials triggered when the rat was in B sectors, the reward size was three times larger (this was to encourage rats to use B sectors; see below). In trials triggered when the rat was in C sectors, the reward size was randomly selected to be 1–4 times that of an A sector.

### MDP formulation of the rat task

#### General setup

For the simplest model, the position of the rat was discretized into 19 logically defined positions as shown in Fig. 1B, named A1–A6, B1–B6, C1–C6, and D. MDP states are defined as tuples *(position, IA-curr, IA-prev, trial-type)*, where *IA-curr* denotes a currently active interaction area, and *IA-prev* the previously active interaction area. Both *IA-curr* and *IA-prev* are a number between zero and six, where zero indicates that no interaction area is active, and positive numbers refer to the interaction area of the same number. The *trial-type* is either A, B, C, or P, where A, B, C indicate that the upcoming reward was triggered from an A, B, C area respectively, and P indicates that the current trial is a warning trial. This model has 19 × 7 × 7 × 4 = 3724 states.

MDP actions were formalized as follows. One movement action was defined for each of the 19 defined sectors of the MDP. When selecting actions, movement actions were available only between neighboring MDP areas. For example, in position B3, only movements A3, C2, C3, and D were available. In addition to the 19 movement actions, two poke actions were defined, “FD” for the food port and “WA” for the water port of an interaction area. An activate action “AC” was defined to indicate activation of an interaction area; this action corresponded to the TGT event described above. Based on the rat’s position during the “AC” action, an IA became active, and the *trial-type* and *IA-curr* of the next MDP state were set accordingly. Timeouts were coded by a timeout action “TO”, used whenever an interaction area became inactive due to the rat’s inactivity.

MDP rewards were set to +30 for both water and food. Airpuffs were assigned a reward of -30. Each movement was assigned a cost in the following way: movements between A and neighboring B areas were assigned a reward of -1, between A and neighboring C areas the reward was -1.5, and between B and D it was -1.

In our task, most state transitions were deterministic given the current state and the selected action. The following were exceptions: when no IA is active and the rat selected the “activate” action in an A, B or C sector (i.e., the rat waited in that sector), an IA was activated with probability 0.9, and a “warning” trial was initiated with probability 0.1. In the case of A and B sectors, the activated IA is determined deterministically, while in a C sector, one out of six IAs is selected for activation with uniform probability.

Decoding the observed rat trajectories into a sequence of MDP states and their transitions was performed in the natural way. Rat movements and nose-pokes were directly translated into the corresponding MDP actions. State transitions that were triggered by waiting for an interaction area to become active were translated to the “AC” action, and timeouts to the “TO” action.

#### The augmented MDP

We extended the MDP model to include costs for sharp-angled body rotations by discretizing the rat body orientation into four values relative to the arena: outward-looking, clockwise-looking, inward-looking, and counter-clockwise-looking (in Fig. 1B, the magenta/orange/blue rats are outward-/clockwise-/inward looking, respectively). In this MDP, states were coded as *(position, body-orientation, IA-curr, IA-prev, trial-type).* The body orientation was updated as a consequence of movement. For instance, moving from a B sector to an adjacent A sector entails an outward-looking body orientation.

We created families of MDPs parametrized by body rotation cost multiplier *m*_orientation_ and poke reward *r*_poke_. These MDPs had the same set of states and actions, as well as the same state transition matrix. However, their reward functions differed. The body rotation costs were given by −*m*_orientation_ for a 90 degree turn, and −2 *× m*_orientation_ for a 180 degree turn. These costs added to those of the movements implied by that state transition. In addition, the rewards for the nose-poke actions (food and water) were increased by *r*_poke_, independently of the status of the IA that was accessed. Consequently, even pokes into non-target ports provided reward. We created a total of 150 MDPs, for all possible combinations of *m*_orientation_ ∈ [0, 1, 2, 3, 4, 5, 6, 7, 8, 10, 12, 15, 20, 25, 30] and *r*_poke_ ∈ [0, 2, 4, 6, 8, 10, 15, 20, 25, 30].

### Computing information-constrained policies

Stochastic policies were generated via a custom Matlab implementation of G learning (Fox, Pakman, and Tishby 2016). G learning computes optimal policies under information constraints. In particular, for a policy *π* and a state *s*, define the value of *s* as

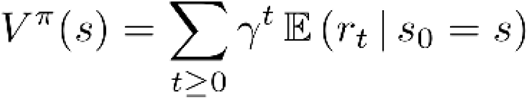

where the expectation is taken over all trajectories with starting state *s*, *r_t_* is the reward at time step *t*, and *γ* is a discount factor, slightly smaller than one.

We defined the expected informational cost of *π* at starting state *s* as

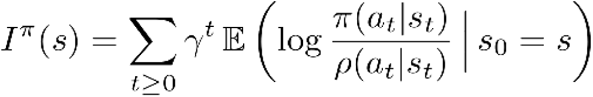

where *ρ* is a task-agnostic policy, in our case the policy with equal probabilities for all accessible actions at each state. The information cost measures the deviation of *π* from the default policy *ρ* (for details see Fox, Pakman, and Tishby (2016)).

G learning is an algorithm for computing policies *π* that minimize the *free energy*

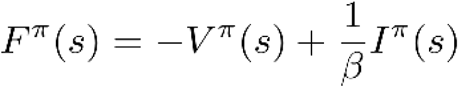

for all states simultaneously. The free energy contains a single parameter *β* that controls the trade-off between value (future total reward/cost in the MDP) and information cost. At low *β*, the information cost dominates and forces the policy *π* to be similar to the default policy. At high *β*, the information cost is negligible and *π* approximates an optimal policy.

In all computations, we used a fixed discount factor *γ* = 0.99. We employed G learning by iterating over lists of precomputed MDP states and actions. Each such list consisted of 128 352 tuples *(state_in_, action, state_out_, reward)*, chosen such that each possible combination of *state_in_, action, state_out_, and reward* occurred in the list with the frequency prescribed by the MDP. For instance, as the “activate” action initiates a “warning” trial with probability 0.1, tuples corresponding to activating a “warning” trial were represented with a frequency of 0.1 among all relevant “activate” actions. The input to each epoch of G learning was the list of tuples described above, in random order. We used a dynamic learning rate

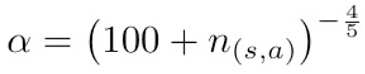

where *n_(s, a)_* counts occurrences of the state-action pair *(s, a)* across epochs of G learning. For each parameter set, G learning was stopped after 1000 epochs or when the largest update of any policy entry between iterations dropped below 0.0001. In practice, around 100 epochs sufficed for small values of *β*, and the full 1000 epochs were needed only for the few largest *β*.

Policies were generated for 200 values of *β*, logarithmically spaced between 0.005 to 1.1. This range was determined by numerical experiments as the range where policies transitioned between being essentially equal to the default policy to being essentially equal to the optimal policy. In total, we computed 30 000 policies (resulting from all combinations of 15 orientation costs, 10 nose-poke rewards, and 200 *β* values).

To obtain an estimated upper bound on performance in the task, we used the policy with the highest value of *β* (i.e., *β* = 1.1). We verified that further increasing *β* did not yield policies of higher value, confirming the optimality of this policy.

### Data analysis

For each experimental session, data were acquired and processed as described previously (Jankowski, Polterovich, et al. 2023). In particular, behavioral data from each session consisted of rat locations, sound presentations, rat nose-pokes, and reward and air-puff deliveries. For each session, data were checked for integrity and converted to a sequence of MDP states and actions, denoted as *(s_t_, a_t_)_t = 1, …, N_*, where *N* denotes the number of state-action pairs observed in the session.

#### Best fitting policies

For a given policy *π_j_*, the log likelihood of a trajectory under *π_j_* is given by

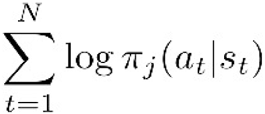

Thus, for a family of policies *(π_j_)_j=1, …, K_*, a best-fitting policy is obtained by maximizing the log likelihood:

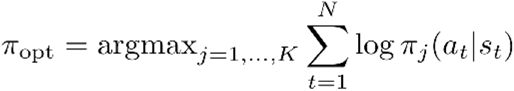

Our policies are parameterized by *β* and in some analyses also by body orientation cost and nose-poke reward. The best-fitting values for these parameters were obtained as the parameters of that policy that had the maximum likelihood with respect to the observed data.

To investigate the evolution of *β* during training, likelihood maximization was performed for each behavioral session using 200 policies with varying *β* computed for the MDP with body orientation cost and poke-reward fixed at zero (different orientation costs and poke-rewards define different MDPs, so it is not meaningful to compare *β* across models where orientation costs and poke-rewards vary). Sample log likelihood functions for two sessions, as a function of *β*, are shown in Fig. 2D. To investigate the evolution of best-fitting orientation costs and poke-rewards, likelihood maximization was performed over the 30 000 policies that were computed also for the MDPs with varying orientation costs and poke rewards. An example of the dependence of the log likelihood on the cost parameters is shown in Fig. 3D.

#### Simulations

Given a policy *π*, the behavior of agents following *π* can be simulated by drawing actions according to *π* and state transitions and rewards according to the underlying MDP. Four policies were compared with the observed data. Two of these policies, the best-fitting 3-parameter policy (*β* and two cost parameters) and the 1-parameter best-fitting policy (*β* with the cost parameters set to 0), were individually optimized for each behavioral session by likelihood maximization as described above. The other two policies were the high-*β* policy, where the highest *β* value in our set of policies was used (essentially equivalent to the use of an optimal policy), and the uniform policy.

For each behavioral session (N = 607 sessions, 155 777 trials total) and each model, 160 simulated trajectories consisting of 1000 MDP actions each were computed. For the 1-parameter and 3-parameter models, the best-fitting parameters for each session were used for the corresponding simulation. From these trajectories, two simple behavioral statistics were obtained: (1) the average number of actions needed to conclude a trial (counting all actions after one feedback event up to the next feedback event) and (2) the average success rate. One-way ANOVA and t-tests were used to compare the actions per trial and success rate between the models and with the observed data.

To analyze differences in the distribution of the number of A, B, and C trials in the simulations and in the observed data, the fractions of selected trial types were extracted from the observed data and from the simulations. The Kullback-Leibler divergence (D_KL_) between the trial type distribution in observed data and each model was computed. For statistical analysis, the ANOVA modeled the dependent variable D_KL_ as a function of the four policies. Post-hoc comparisons between the different models were performed by t-tests.

### Rat experiments and electrophysiology

For a complete description of the experimental procedures, see (Jankowski, Polterovich, et al. 2023). All animal experiments complied with the regulations of the ethics committee of the Hebrew University of Jerusalem. The Hebrew University of Jerusalem is an Association for Assessment and Accreditation of Laboratory Animal Care (AAALAC) accredited institution. Five adult female Sabra rats weighing at least 200 g were used (Envigo LTD, Israel). Animals were food restricted to not lower than 80% their ad-libitum weight. They were trained in the RIFF for five days every week, and had free access to food and water on weekends. Before the main experiment, animals were habituated to the experimenter for up to ten days.

Silicon probes (ASSY-116_E-2, Cambridge Neurotech, UK) were implanted chronically into the left insular cortex. The following coordinates were used as electrode entry point: AP = -1.0 mm, ML = -6.1 mm, relative to Bregma. In a first surgery, a base was prepared for subsequent electrode implantation. In a second surgery (14 to 28 days later), a single axis micromanipulator was used to insert electrodes through a craniotomy performed by drilling.

Training progressed with increasing difficulty, and started with a long habituation period. In order to encourage rats to use all available rewarded strategies, and to gradually introduce “warning” trials into the experiment, the following scheme was used, in all rats, before electrode implantation:

**Methods Table:** Experiment phases before electrode implantation.

| Phase | Weeks | A reward | B reward | Warning probability |
| --- | --- | --- | --- | --- |
| 1 | 1 - 4 | 1 pellet | 1 pellet | 0 |
| 2 | 5 - 7 | 0 pellets | 3 pellets | 0 |
| 3 | 8 - 10 | 1 pellet | 3 pellets | 0 |
| 4 | 11 - 12 | 1 pellet | 3 pellets | increasing from |
|  |  |  |  | 0 to 0.1 |
| 5 | 13 - 18 | 1 pellet | 3 pellets | 0.1 |

After the 18 weeks outlined in Table 1, the five rats were split into three groups: two rats were immediately implanted with electrodes and continued training after recovery, and two rats were implanted with electrodes 1 year (362 days) after the start of Phase 1. The fifth rat was implanted 1.5 years (547 days) after the start of Phase 1, but we did not analyze the electrophysiological data from that rat. Data analyzed in this study is from Phase 1 above, and from the entire period from electrode implantation until the end of the experimental sessions.

