## Supplemental Figures and Tables for "Insular cortex encodes task alignment"

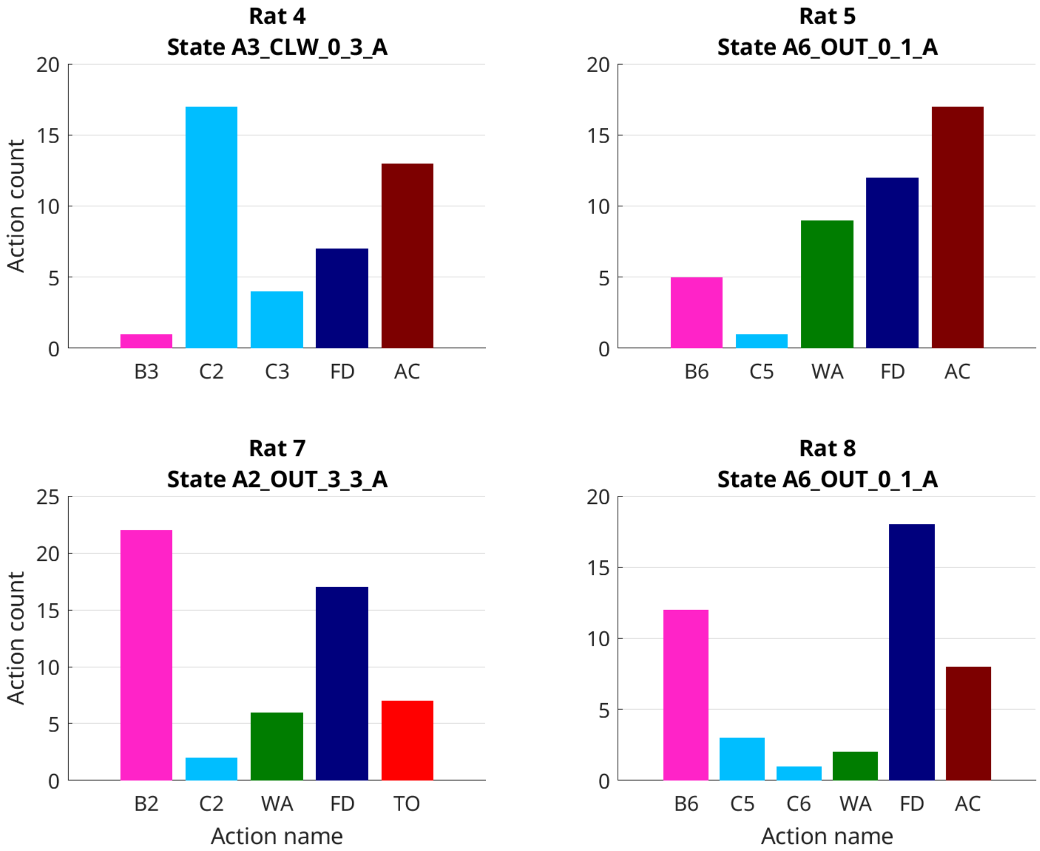


Supp. Fig. 1. **Examples of non-deterministic rat behavior.** Shown are examples from one session per rat, for four different rats. In each panel, a state was selected that occurred a few dozens of times during the session, and in which the rat showed highly non-deterministic behavior.

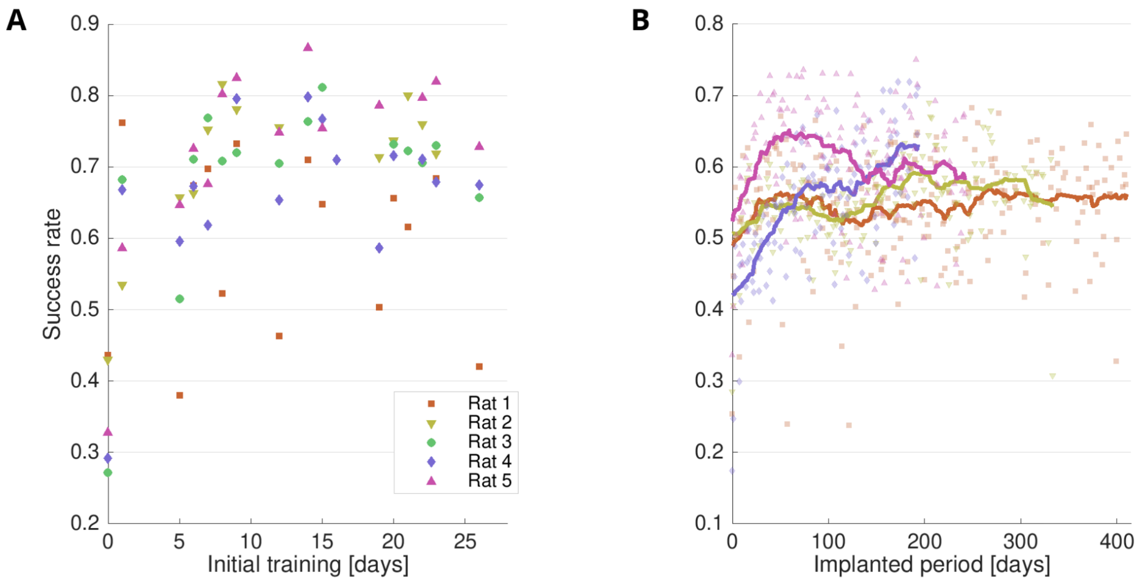


Supp. Fig. 2. **Evolution of rat success rates over time.** A, initial month of training. B, time period after electrode implantation.

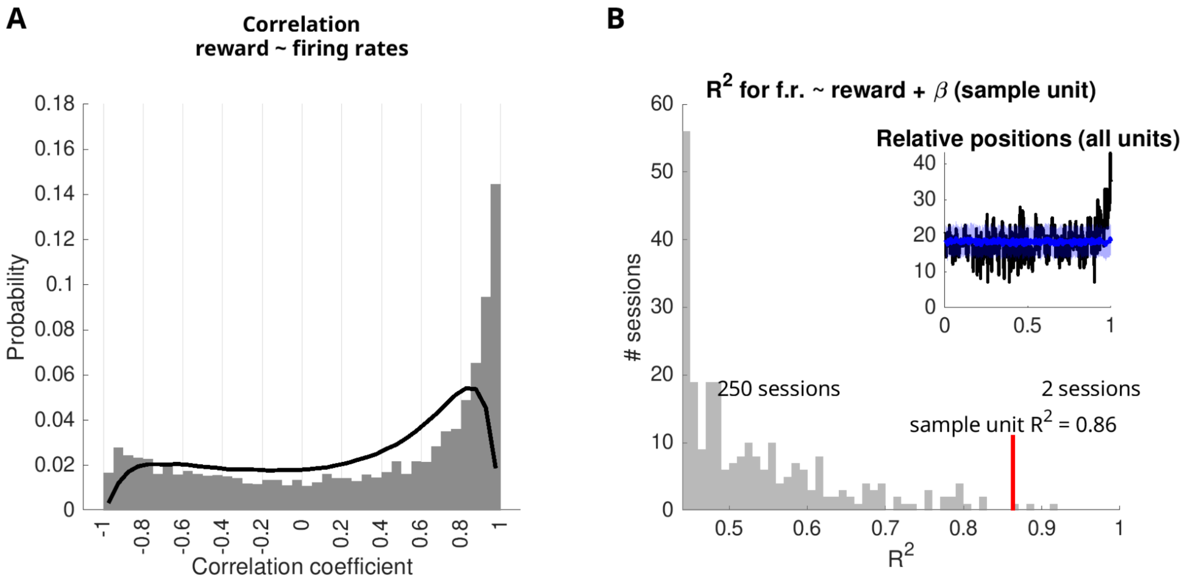


**Supp. Fig. 3. Insular cortex neurons are correlated with task alignment also when controlling for reward events.**

(A) Reward events are correlated with firing rates. The histogram shows Pearson correlation coefficients for reward rates and firing rates in 10-minute windows, where reward rates and firing rates originate from the same session. The continuous line shows the control condition, correlation coefficients for reward rates and firing rates from different sessions.

(B) Firing rates in many neurons correlate more strongly with the same session's reward and β, than with the same session's reward and another session's β. A sample unit is shown in the main plot. The red line indicates the $R^2$ for this unit's firing rate modeled linearly with variables reward and β. The histogram shows $R^2$ for the models of the firing rate of that unit with the same-session reward but β from all 252 other sessions. The inset shows a histogram of all units' relative position for 'same session $R^2$' compared to 'other session $R^2$'. Black line, original data. Blue line, relative positions for randomly permuted session numbers. Blue shading, standard deviation across 100 session number permutations.

| Estimate | F | DF1 | DF2 | p | Time after day | Variable |
|---|---|---|---|---|---|---|
| 0.00642 | 13.7 | 1 | 1274 | 2.3E-04 | 0 | TA |
| 0.00393 | 82.4 | 1 | 1036 | 5.6E-19 | 50 | TA |
| 0.00317 | 10.7 | 1 | 586 | 1.1E-03 | 150 | TA |
| -0.00412 | 72.7 | 1 | 1274 | 4.3E-17 | 0 | Interaction |
| -0.00450 | 59.3 | 1 | 1036 | 3.2E-14 | 50 | Interaction |
| -0.00525 | 31.3 | 1 | 586 | 3.4E-08 | 150 | Interaction |

**Supp. Table 1. Estimates of the change in success rates and task alignment for three different time periods.** Estimated is the daily change in standardized units. The change for success rates is obtained as the sum of the changes for task alignment and the interaction.

| | Actions per trial | | | | Success rate | | | |
|---|---|---|---|---|---|---|---|---|
| **Model** | Mean | SEM | Min | Max | Mean | SEM | Min | Max |
| Observed | 5.43 | 0.046 | 2.73 | 13.45 | 0.559 | 0.0033 | 0.174 | 0.75 |
| Three-param. | 6.10 | 0.023 | 4.96 | 9.74 | 0.712 | 0.0029 | 0.279 | 0.88 |
| One-param. | 6.14 | 0.022 | 4.99 | 9.74 | 0.742 | 0.0027 | 0.279 | 0.88 |
| High beta | 3.80 | 0.0005 | 3.77 | 3.84 | 0.999 | 0.0001 | 0.996 | 1.00 |
| Uniform | 10.27 | 0.009 | 9.68 | 10.87 | 0.200 | 0.0005 | 0.164 | 0.24 |

**Supp. Table 2. Behavioral measures "actions per trial" and "success rate" for observed sessions, and for four different models.**

| Model | A frac. | B frac. | C frac. |
|---|---|---|---|
| Observed | 0.50 | 0.26 | 0.13 |
| Three-param. | 0.40 | 0.38 | 0.12 |
| One-param. | 0.32 | 0.45 | 0.13 |
| High beta | 0.01 | 0.89 | 0.00 |
| Uniform | 0.33 | 0.28 | 0.29 |

**Supp. Table 3. Average selection of A, B, and C trials for observed sessions, and for four different models.** Difference to 1 is due to 10% "warning" trials occurring in each session that did not depend on rats' decisions.
